# Tensor-Derived Similarity Networks for Characterising Spatial Patterns in Colorectal Cancer

**DOI:** 10.64898/2026.04.20.719742

**Authors:** Tuan D. Pham

## Abstract

Spatial transcriptomics enables the study of gene expression within the spatial context of tissue architecture, offering new opportunities for understanding tumour heterogeneity. This study proposes a tensor-derived similarity network framework for analysing spatial organisation in colorectal cancer. Gene expression data from four patients are represented as spatially structured tensors and decomposed using a low-rank canonical polyadic model to extract latent spatial–molecular features. These features are used to construct similarity networks that characterise spatial relationships between tissue regions. Global network measures, including similarity, density, and spatial heterogeneity, reveal sparse but structured connectivity patterns across all patients. An embedding-permutation framework is introduced to generate randomised spatial configurations while preserving feature distributions. Comparative analysis shows that randomised networks exhibit higher similarity, density, and heterogeneity than real data, indicating that spatial organisation constrains network structure. The results demonstrate that the proposed framework captures meaningful spatial patterns in tumour tissue and provides quantitative measures of spatial heterogeneity. This approach offers a general methodology for analysing spatial transcriptomics data and has potential applications in spatial biomarker discovery and characterisation of tumour architecture.

## 1 Introduction

Spatial transcriptomics has emerged as a powerful technology for measuring gene expression while preserving the spatial context of tissue architecture. Unlike conventional bulk or single-cell sequencing approaches, spatial transcriptomics enables the study of molecular activity within its native spatial environment, providing new opportunities to investigate tissue organisation, cellular interactions, and tumour heterogeneity [1, 2, 3, 4, 5, 6, 7, 8]. In colorectal cancer (CRC), spatial heterogeneity is a defining feature of tumour biology, influencing tumour progression, immune response, and therapeutic outcome [9, 10, 11, 12, 13].

The tumour microenvironment in CRC is highly complex, consisting of malignant epithelial cells, stromal components, immune infiltrates, and vascular structures arranged in spatially organised niches [14, 15, 16]. These niches include regions of immune activation, immune suppression, hypoxia, and stromal invasion, each characterised by distinct gene expression profiles. The spatial arrangement of these regions plays a critical role in determining tumour behaviour. For example, the presence and localisation of tumour-infiltrating lymphocytes have been associated with improved prognosis, while spatial exclusion of immune cells or the formation of immunosuppressive niches can contribute to tumour progression and resistance to therapy [17, 18, 19]. Consequently, understanding not only the molecular composition but also the spatial organisation of tumour tissue is essential for advancing precision oncology in CRC.

Beyond CRC, spatial biology has broad applications across multiple domains of medicine and biology. In oncology, spatial transcriptomics has been applied to diverse tumour types, revealing spatially distinct tumour subclones, immune niches, and stromal interactions [20, 21, 22, 23]. In immunology, spatial analysis enables identification of functional immune microenvironments, which are critical for understanding responses to immunotherapy [24, 25]. In neuroscience and developmental biology, spatial gene expression patterns provide insights into tissue organisation, functional architecture, and developmental processes [26, 27].

Current analytical approaches, including tensor decomposition and graph theory, for spatial transcriptomics data typically rely on clustering, spatial statistics, cell-type deconvolution, multi-scale dynamics, or image-based segmentation to identify regions of interest [28, 29, 30, 31, 32, 33, 34]. While these methods can detect spatial domains or gradients, they often treat spatial locations independently or rely on predefined structures, limiting their ability to capture complex spatial–molecular interactions. In particular, there remains a need for computational frameworks that can integrate high-dimensional gene expression with spatial organisation in a unified and quantitative manner.

Tensor-based methods [35, 36, 37] provide a natural representation for multi-dimensional data, enabling integration of spatial and molecular information within a single mathematical framework. By organising spatial transcriptomics data into higher-order tensors, latent spatial–molecular patterns can be extracted that reflect underlying biological organisation. Tensor decomposition has also been applied in medical imaging [38, 39, 40], genomics [41], and multiomic data [42, 43]. However, tensor representations alone do not directly quantify interactions between spatial regions or provide interpretable measures of tissue organisation. Network-based approaches offer a complementary perspective by modelling relationships between spatial locations as graphs, where nodes represent spatial spots and edges encode similarity or connectivity. In systems biology, networks have been extensively used to study gene regulation, protein interactions, and signalling pathways [44, 45, 46, 47, 48]. Extending this paradigm to spatial networks enables the investigation of how biological interactions are organised within physical space. However, network-based approaches typically lack direct integration with high-dimensional molecular representations.

These limitations motivate the development of computational frameworks that combine the strengths of both tensor-based modelling and network analysis. In particular, there is a need for methods that can jointly capture high-dimensional molecular variation, spatial organisation, and interaction structure in a coherent and interpretable manner. Furthermore, distinguishing biologically meaningful spatial patterns from those arising due to random variation remains a key challenge in spatial transcriptomics analysis.

In this study, a tensor-derived similarity network framework is developed for analysing spatial transcriptomics data in CRC. Spatial gene expression is first represented as a tensor and decomposed using a low-rank canonical polyadic model to extract latent spatial–molecular features. This representation enables the integration of high-dimensional gene expression with spatial organisation, providing a structured description of tissue architecture. The resulting latent features are subsequently used to construct similarity networks that characterise relationships between spatial regions, allowing interactions between tissue locations to be analysed in a graph-based setting. Network-based measures are introduced to quantify global similarity, connectivity, and spatial heterogeneity, thereby providing interpretable descriptors of spatial organisation.

By linking tensor decomposition with network modelling, the proposed framework establishes a unified approach for capturing both latent spatial–molecular patterns and their interactions. This integration enables the transition from abstract feature representations to measurable properties of tissue organisation, facilitating the quantitative analysis of spatial structure. In doing so, the framework provides a coherent methodological basis for investigating spatial heterogeneity and supports further exploration of spatial transcriptomic patterns in biological and clinical contexts.

## 2 Materials and Methods

### 2.1 Visium Spatial Transcriptomics Dataset

Spatial transcriptomics data were obtained from the publicly available dataset [49] “Visium spatial transcriptomics analysis of human primary colorectal cancer” (GEO accession: GSE226997). This dataset comprises spatially resolved gene expression measurements generated using the 10x Genomics Visium Spatial Gene Expression platform from human primary CRC tissue sections. The data provide spatially localised transcriptomic information, enabling the investigation of tumour heterogeneity and tumour microenvironment organisation.

The Visium platform captures gene expression across tissue sections using spatially barcoded capture spots arranged on a predefined grid. Each capture spot contains spatial barcodes that allow transcriptomic measurements to be mapped back to their physical tissue locations. In the GSE226997 dataset, spatial transcriptomics measurements were generated from four human primary CRC specimens, providing independent samples for cross-patient comparison of tumour architecture and spatial heterogeneity.

For each tissue section, the dataset includes a spatially resolved gene expression matrix, spot barcodes, gene annotations, spatial coordinates, and histological tissue images. Gene expression data are stored in the file `filtered_feature_bc_matrix.h5`, which contains filtered count data following initial quality control using the 10x Genomics Space Ranger pipeline. The matrix is represented in sparse format using non-zero expression values, row indices corresponding to genes, and column pointer indices defining the sparse structure.

Spatial coordinates for each capture spot are provided in the file `tissue_positions_list.csv`, which includes spot barcodes, tissue indicators, array coordinates, and pixel-level spatial positions. These coordinates enable mapping of gene expression measurements to their physical tissue locations and support spatial analysis of tumour heterogeneity.

The Visium platform typically captures approximately 4,000–5,000 spatial spots per tissue section, with each spot representing gene expression from multiple cells within a capture diameter of approximately 55 *µ*m. In the GSE226997 dataset, 4,148 spatial spots and 17,944 genes were retained following quality control, providing comprehensive spatial gene expression coverage of colorectal tumour tissue.

Additional preprocessing was performed prior to analysis. Spatial spots outside tissue regions were removed using the in tissue indicator. Spots with zero or extremely low gene expression counts were excluded to ensure data quality. Gene expression values were normalised using library-size normalisation followed by logarithmic transformation:

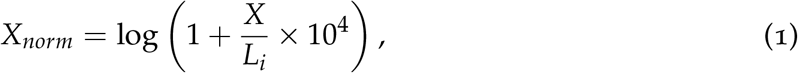

where 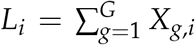 denotes the library size for spatial spot *i*, defined as the total gene expression counts across all genes. This normalisation reduces technical variability arising from differences in sequencing depth and RNA capture efficiency.

Highly variable genes were subsequently selected to retain biologically informative features while reducing noise. Principal component analysis (PCA) was then applied to obtain low-dimensional molecular representations suitable for downstream tensor construction and similarity network analysis.

The four CRC samples in GSE226997 provide independent spatial transcriptomics measurements, enabling cross-patient comparison of tumour architecture and assessment of reproducibility of spatial heterogeneity patterns. The processed spatial transcriptomics data were subsequently used for tensor construction, tensor decomposition, and similarity network analysis. Figure 2 shows brightfield histological images of spatial transcriptomics tissue sections from four CRC patients obtained from the GSE226997 dataset. These images illustrate the heterogeneous tumour morphology, glandular organization, stromal composition, and microenvironmental variability across patients. The overlaid capture array grid indicates spatial measurement locations, where each spot corresponds to a spatially resolved gene expression profile. These patient-specific tissue architectures provide the morphological foundation for subsequent tensor construction and spatial similarity network analysis to quantify tumour heterogeneity across patients.

**Figure 1:**
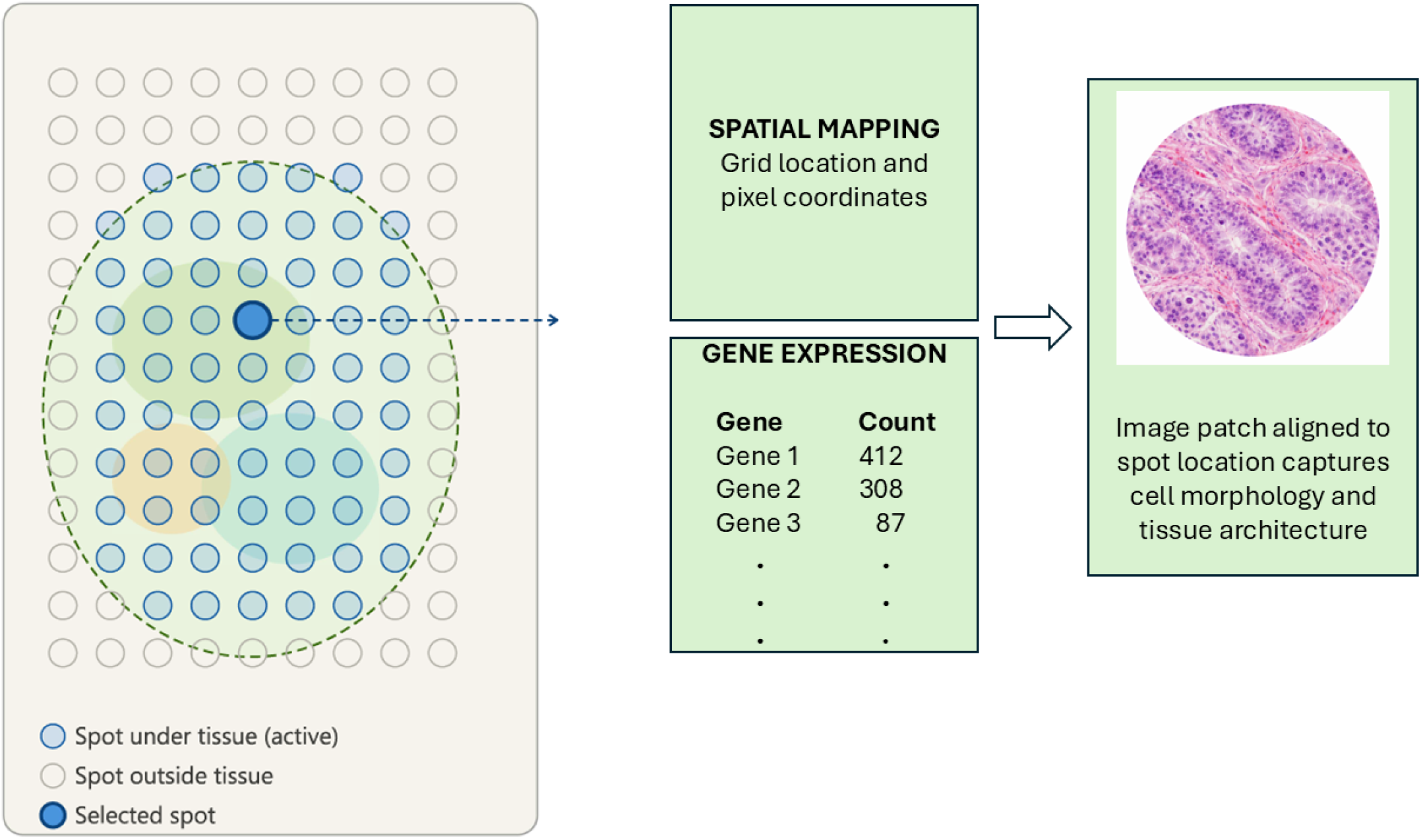
Schematic of spatial transcriptomics data structure. A rectangular capture array of spots is printed on a slide, and a tissue section is placed on top. Spots beneath the tissue (filled circles) capture messenger RNA from the cells above them, while spots outside the tissue boundary (hollow circles) remain inactive. (B) Each active spot yields three linked data components: (i) a pair of spatial coordinates recording its row and column position within the array, which preserves its location in the tissue; (ii) a gene expression count vector recording the number of transcripts detected per gene across thousands of genes; and iii) a histological image patch derived from brightfield imaging of the tissue, capturing local cell morphology and tissue architecture co-registered to the spot location. Together, these components enable gene expression to be mapped and analysed in its spatial tissue context.

**Figure 2:**
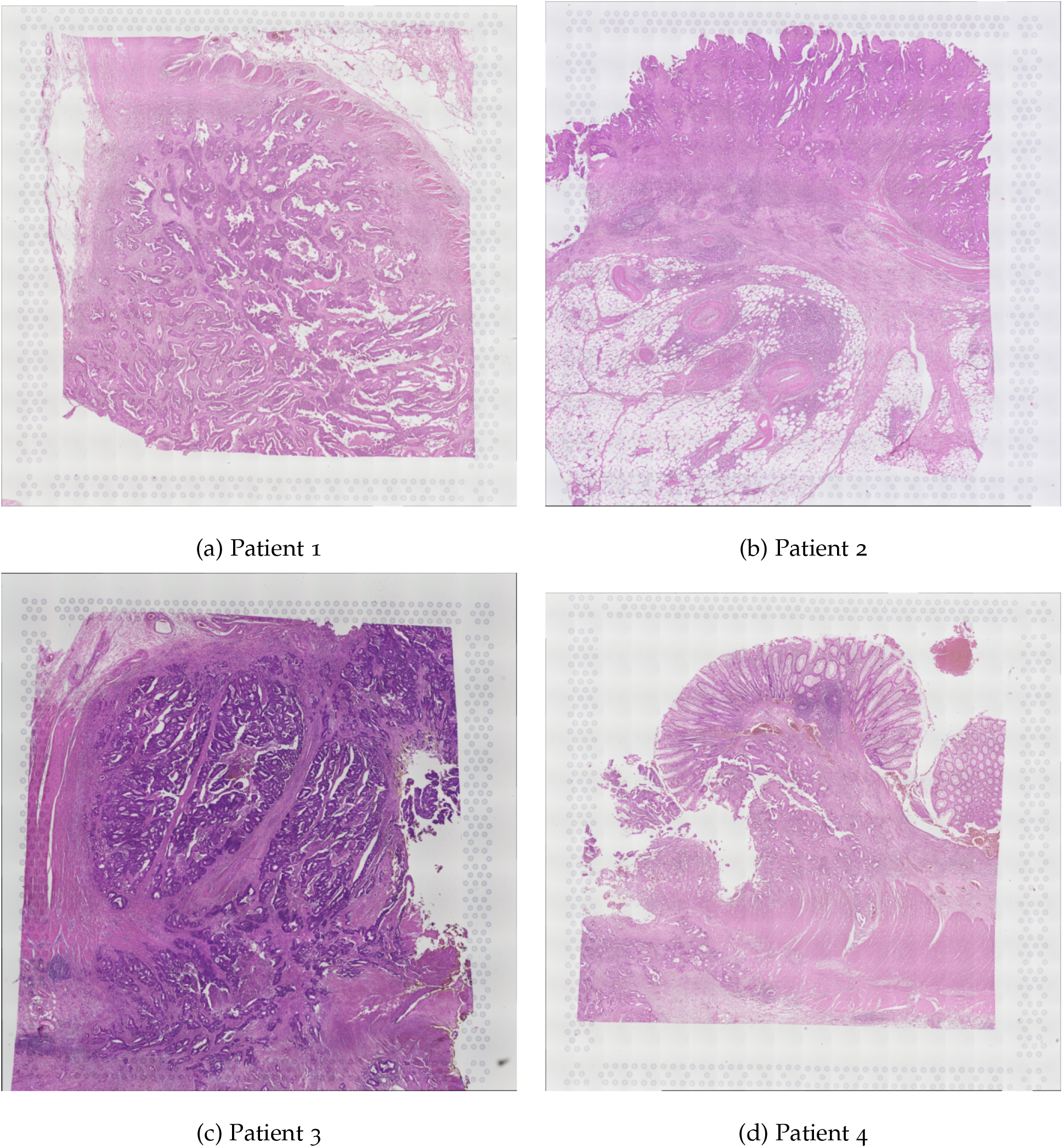
Spatial transcriptomics histological images (low resolution) from GSE226997 colorectal cancer dataset. Brightfield hematoxylin and eosin (H&E) stained tissue section acquired using the 10x Genomics Visium platform. The faint circular grid surrounding the tissue corresponds to the Visium capture array, where each spot represents a spatial location associated with gene expression measurements. Spots located beneath the tissue capture transcriptomic information, while spots outside the tissue remain inactive.

**Figure 3:**
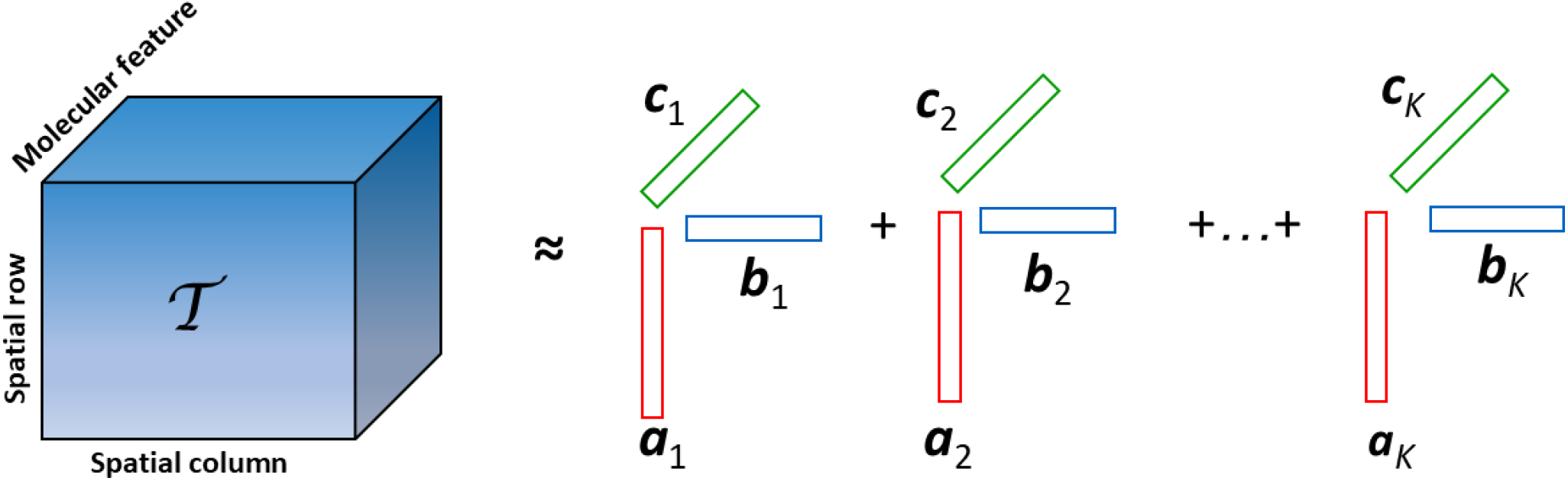
Canonical polyadic decomposition of a three-way tensor into rank-one components.

### 2.2 Tensor Decomposition and Similarity Networks for Spatial Tumour Heterogeneity

Let *G* denote the number of genes and *N* denote the number of spatial spots. The spatial transcriptomics dataset is represented as a gene expression matrix

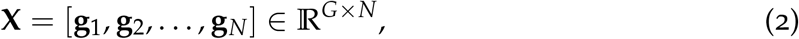

where **g**_*i*_ ∈ ℝ^*G*^ denotes the gene expression vector at spatial location *i*. Each spatial spot is associated with spatial coordinates (*x*_*i*_, *y*_*i*_) describing its physical location within the tissue.

#### Dimensionality reduction

To reduce the high dimensionality of gene expression data, principal component analysis (PCA) is applied. Let

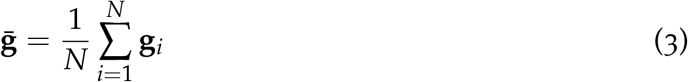

denote the mean gene expression vector, and define the centred data matrix

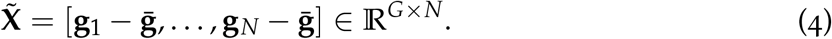

The covariance matrix is given by

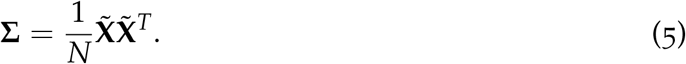

Eigenvalue decomposition yields

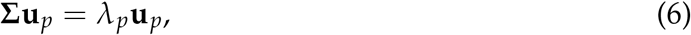

where *λ*_*p*_ and **u**_*p*_ denote eigenvalues and eigenvectors, respectively. The principal component matrix is

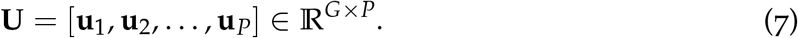

The reduced feature matrix is obtained as

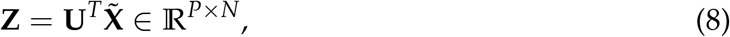

where the *i*-th spatial spot is represented by

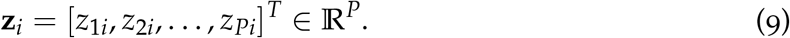

#### Tensor construction

To integrate spatial and molecular information, a third-order tensor is constructed:

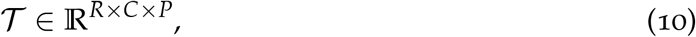

where *R* and *C* denote the spatial grid dimensions. Let (*r*_*i*_, *c*_*i*_) denote the grid location corresponding to spatial spot *i*. The tensor entries are defined as

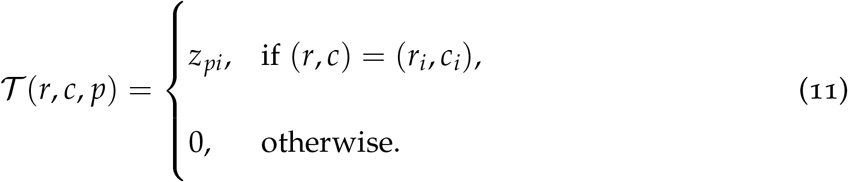

#### Tensor decomposition

The tensor is decomposed using canonical polyadic (CP) decomposition or PARAFAC (parallel factor analysis) [36, 50],:

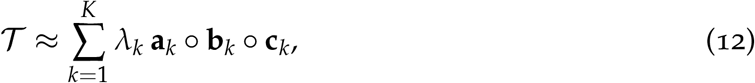

where *λ*_*k*_ denotes the weight of the *k*-th component, **a**_*k*_ ∈ ℝ^*R*^ and **b**_*k*_ ∈ ℝ^*C*^ represent spatial factors, and **c**_*k*_ ∈ ℝ^*P*^ represents molecular feature factors.

Element-wise, the decomposition can be written as

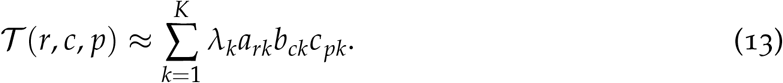

#### Tensor-derived embedding

The spatial activation of component *k* at spot *i* is defined as

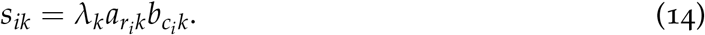

Each spatial spot is embedded in a *K*-dimensional latent space:

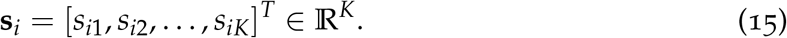

#### Similarity network construction

Similarity between spatial spots is computed as

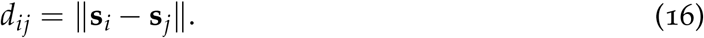

A Gaussian kernel defines the similarity matrix:

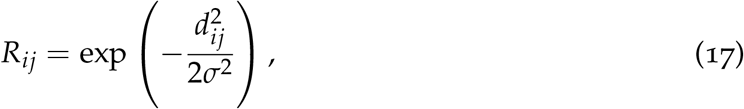

where *σ* = median_*i*<*j*_(*d*_*ij*_).

#### Spatial constraint

Let **v**_*i*_ = (*x*_*i*_, *y*_*i*_). The *k*-nearest neighbours are

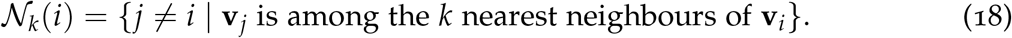

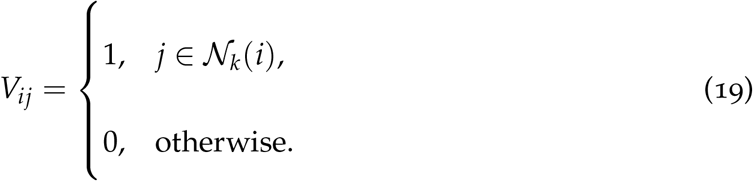

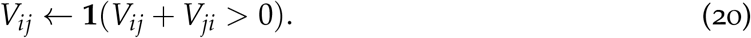

where **1**(·) denotes the indicator function.

The spatially constrained similarity matrix is defined as

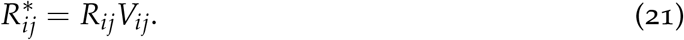

#### Global similarity

The global similarity rate is defined as

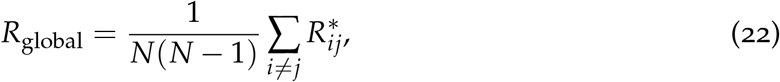

which quantifies the overall level of similarity between spatial spots in the tissue, as captured by the tensor-derived similarity network under spatial constraints. Higher values of *R*_global_ indicate stronger and more widespread similarity among spatial regions, reflecting more homogeneous spatial organisation, whereas lower values suggest greater variability and increased spatial heterogeneity in tumour structure.

#### Network topology

Node strength is defined as

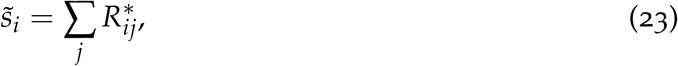

which measures the total similarity between spatial spot *i* and its neighbouring spots in the network. Higher values of 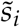 indicate that a spot is strongly connected to similar surrounding regions, reflecting locally homogeneous tissue structure, whereas lower values suggest weaker similarity and greater local spatial variability.

Node degree is defined as

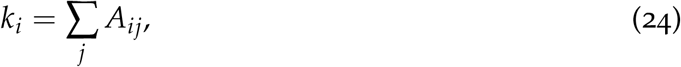

which represents the number of spatial neighbours connected to spot *i* in the thresholded similarity network. Higher values of *k*_*i*_ indicate that a spot shares strong similarity with many neighbouring regions, reflecting locally dense connectivity, whereas lower values suggest fewer similar neighbours and more spatially distinct local structure.

Node density is given by

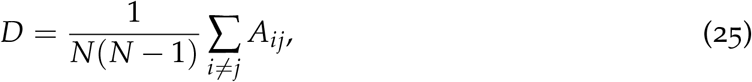

which quantifies the proportion of connections present in the thresholded similarity network relative to all possible connections. Higher values of *D* indicate a more densely connected network with widespread similarity among spatial spots, whereas lower values reflect sparser connectivity and more spatially heterogeneous organisation.

#### Spatial heterogeneity

Local spatial heterogeneity is quantified using Shannon entropy:

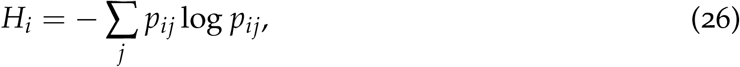

in which

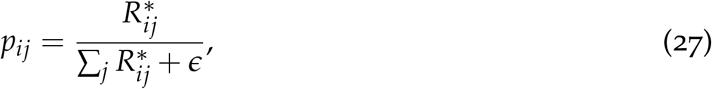

where *ϵ* > 0 is a small constant for numerical stability.

Local spatial heterogeneity *H*_*i*_ quantifies the diversity of similarity interactions between spatial spot *i* and its neighbouring regions. Higher values of *H*_*i*_ indicate a more heterogeneous local environment with a wide range of similarity strengths, reflecting complex and varied tissue structure, whereas lower values correspond to more uniform neighbourhoods with consistent similarity patterns.

Global spatial heterogeneity is defined as

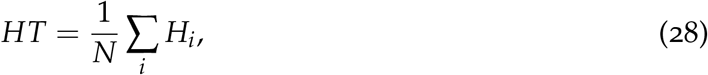

which represents the average level of local heterogeneity across all spatial spots in the tissue. Higher values of *HT* indicate greater overall variability in spatial organisation and more complex tumour architecture, whereas lower values reflect more homogeneous spatial structure.

### 2.3 Embedding-Permutation Framework for Comparative Analysis of Spatial Network Structure

To investigate differences between observed and randomised spatial organisation, an embedding-permutation framework was employed to enable direct comparison between tensor-derived similarity networks and their randomised counterparts. The framework preserves the statistical properties of the learned feature representation while disrupting spatial alignment, thereby isolating the contribution of spatial organisation to network structure.

Following tensor decomposition, each spatial spot is represented by a low-dimensional embedding vector **s**_*i*_ ∈ ℝ^*K*^. Collecting all embeddings yields the matrix

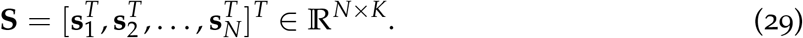

Randomised spatial configurations are generated by permuting the rows of the embedding matrix across spatial locations. For each realisation, a permuted embedding matrix is defined as

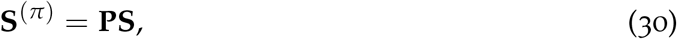

where **P** ∈ ℝ^*N×N*^ is a permutation matrix. This transformation preserves the empirical distribution of tensor-derived features while removing their spatial correspondence.

For each permuted configuration, the spatial similarity network is reconstructed using the same spatial *k*-nearest neighbour graph and Gaussian similarity kernel as defined previously. This ensures that differences between real and randomised networks arise solely from the spatial arrangement of the embedding vectors.

Network measures are then computed for each realisation, including global similarity rate *R*_global_, network density *D*, node strength 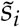, and spatial heterogeneity *HT*. Repeating this procedure over *P* independent embedding-permutation realisations yields a set of network metrics

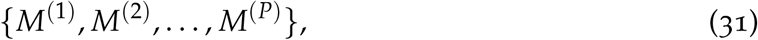

where *M* denotes any of the considered global measures.

From these, summary statistics are computed, including the mean and standard deviation:

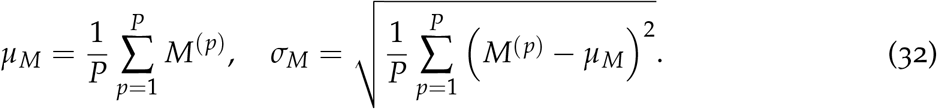

These statistics characterise the typical behaviour and variability of network measures under random spatial organisation. The corresponding metrics computed from the real spatial configuration are compared directly against these values.

This embedding-permutation framework enables systematic evaluation of how spatial organisation influences network topology. Differences in connectivity patterns, network density, and entropy-based heterogeneity between real and randomised configurations characterise the extent to which tensor-derived features capture spatial patterns in the data.

### 2.4 Implementation and Parameter Settings

All analyses were performed using a consistent set of parameters to ensure comparability across datasets. Only spatial spots annotated as tissue were included in the analysis in order to restrict computations to biologically relevant regions. Gene expression dimensionality was reduced by selecting the top 1000 highly variable genes, capturing the most informative transcriptional variation. PCA was then applied, retaining the first 10 components to obtain a compact low-dimensional representation. Missing values were assigned a value of zero to maintain numerical stability and ensure compatibility with downstream tensor construction.

The reduced feature representation was organised into a spatial tensor and decomposed using a CP model with rank *K* = 3. This low-rank approximation captures dominant spatial–molecular patterns while limiting model complexity. The resulting tensor-derived embeddings were used to construct similarity networks, where pairwise similarities between spatial locations were computed and subsequently thresholded with *τ* = 0.5 to obtain a sparse adjacency structure. Spatial constraints were incorporated by restricting connections to a *k*-nearest neighbour graph with *k* = 8, defined in the physical coordinate space, thereby preserving local spatial relationships.

To assess the role of spatial organisation, an embedding-permutation framework was employed. Specifically, 50 independent random permutations of the embedding vectors were generated, producing randomised spatial configurations while preserving the distribution of learned features. Network measures were computed for each permutation to characterise the behaviour of the model under random spatial arrangement.

All experiments were conducted with a fixed random seed to ensure reproducibility of the results.

## 3 Results

### 3.1 Tensor-Derived Similarity Networks Across Patients

The proposed tensor-derived similarity network framework was applied to spatial transcriptomics data from four CRC patients. For each patient, gene expression data were organised into spatial tensors according to the underlying tissue grid and subsequently decomposed to obtain latent spatial–molecular representations.

The dimensional characteristics of the datasets and derived representations are summarised in Table 1. The number of spatial spots varied across patients, resulting in tensor dimensions ranging from 72 *×* 125 *×* 10 to 78 *×* 128 *×* 10. The tensor decomposition produced low-dimensional embeddings of size *N ×* 5 for each patient, where *N* denotes the number of spatial spots. These embeddings form the basis for the construction of similarity networks used in subsequent analysis.

**Table 1:**
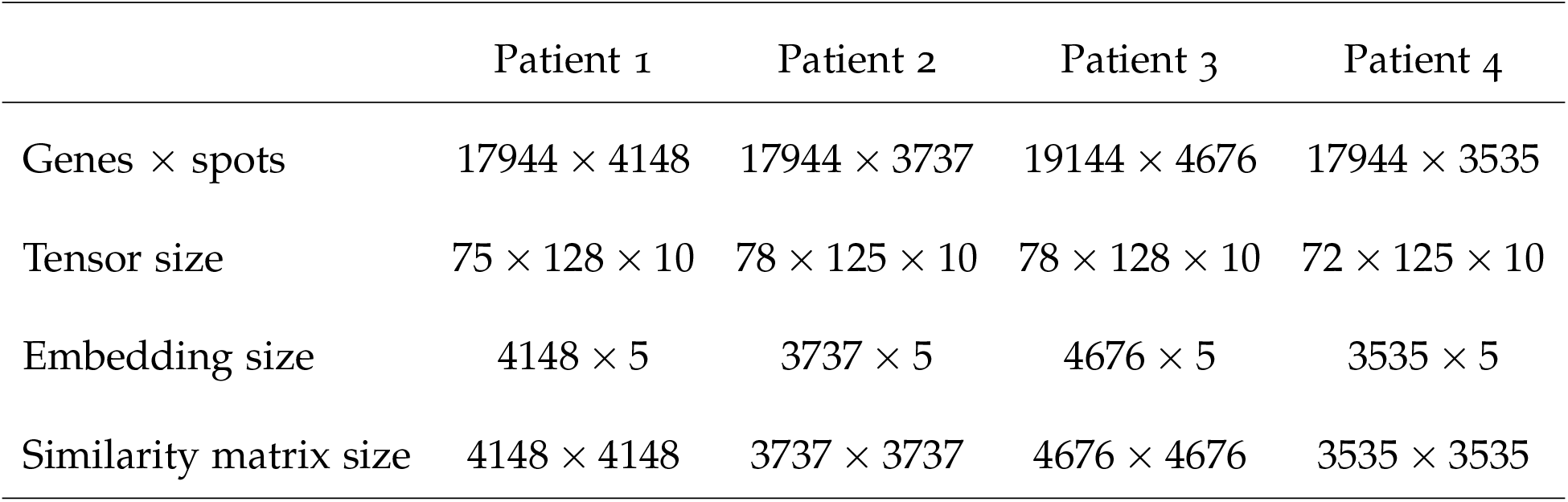
Summary of data dimensions and tensor representations for each patient.

|  | Patient 1 | Patient 2 | Patient 3 | Patient 4 |
| --- | --- | --- | --- | --- |
| Genes × spots | $17944 \times 4148$ | $17944 \times 3737$ | $19144 \times 4676$ | $17944 \times 3535$ |
| Tensor size | $75 \times 128 \times 10$ | $78 \times 125 \times 10$ | $78 \times 128 \times 10$ | $72 \times 125 \times 10$ |
| Embedding size | $4148 \times 5$ | $3737 \times 5$ | $4676 \times 5$ | $3535 \times 5$ |
| Similarity matrix size | $4148 \times 4148$ | $3737 \times 3737$ | $4676 \times 4676$ | $3535 \times 3535$ |

### 3.2 Global Network Measures

Global similarity rate, network density, and spatial heterogeneity were computed from the tensor-derived similarity networks, with results summarised in Table 2. Across all patients, the networks exhibit low values of *R*_global_ and *D*, indicating sparse connectivity. The largest connected component contains the majority of nodes in each case. The spatial heterogeneity measure *HT* varies across patients.

**Table 2:** Global network measures derived from real spatial configurations.

|  | Patient 1 | Patient 2 | Patient 3 | Patient 4 |
| --- | --- | --- | --- | --- |
| Global similarity $R_{\text{global}}$ | 0.000834 | 0.000886 | 0.000958 | 0.001355 |
| Network density $D$ | 0.000692 | 0.000790 | 0.000857 | 0.001400 |
| Global heterogeneity $HT$ | 1.542049 | 1.435756 | 1.705912 | 1.733868 |
| Largest component | 4074 | 3737 | 4119 | 3026 |

### 3.3 Embedding-Permutation Framework

The embedding-permutation framework was applied to generate randomised spatial configurations while preserving the distribution of tensor-derived embeddings. For each patient, multiple independent permutations were performed, and network measures were computed for each realisation.

The mean and standard deviation of the resulting network metrics are summarised in Table 3, characterising the behaviour of similarity networks under random spatial organisation.

**Table 3:** Embedding-permutation network measures (mean *±* standard deviation).

|  | Patient 1 | Patient 2 | Patient 3 | Patient 4 |
| --- | --- | --- | --- | --- |
| $R_{\text{global}}$ | $(2.323 \pm 0.003) \times 10^{-3}$ | $(2.669 \pm 0.004) \times 10^{-3}$ | $(1.964 \pm 0.003) \times 10^{-3}$ | $(2.616 \pm 0.005) \times 10^{-3}$ |
| $D$ | $(2.590 \pm 0.007) \times 10^{-3}$ | $(3.121 \pm 0.010) \times 10^{-3}$ | $(2.109 \pm 0.006) \times 10^{-3}$ | $(2.772 \pm 0.009) \times 10^{-3}$ |
| $HT$ | $2.7093 \pm 5.08 \times 10^{-4}$ | $2.7309 \pm 1.72 \times 10^{-4}$ | $2.6775 \pm 5.52 \times 10^{-4}$ | $2.6826 \pm 6.25 \times 10^{-4}$ |

The reconstructed networks were based on sparse spatial graphs, with the number of undirected edges ranging from 28,213 to 37,343 across patients. Tensor decomposition was applied consistently, with reconstruction errors in the range 0.939940 to 0.944683.

### 3.4 Comparison between Real and Randomised Configurations

A comparison between real and embedding-permuted configurations is summarised in Table 4. The real network measures are reported alongside the mean values obtained from the embedding-permutation framework. Across all patients, the embedding-permutation framework produced networks with consistently higher global similarity rates, higher network densities, and higher spatial heterogeneity compared to the real spatial configurations.

**Table 4:** Comparison of real and embedding-permuted network measures.

|  | Patient 1 | Patient 2 | Patient 3 | Patient 4 |
| --- | --- | --- | --- | --- |
| $R_{\text{global}} (\times 10^{-3})$ | | | | |
| Real | 0.834 | 0.886 | 0.958 | 1.355 |
| Permuted (mean) | 2.323 | 2.669 | 1.964 | 2.616 |
| $D (\times 10^{-3})$ | | | | |
| Real | 0.692 | 0.790 | 0.857 | 1.400 |
| Permuted (mean) | 2.590 | 3.121 | 2.109 | 2.772 |
| $HT$ | | | | |
| Real | 1.542 | 1.436 | 1.706 | 1.734 |
| Permuted (mean) | 2.709 | 2.731 | 2.677 | 2.6826 |

### 3.5 Visualisation of Tensor-Derived Similarity Networks

The tensor-derived similarity network results for the four patients are illustrated in Figures 4, 5, 6, and 7. For each patient, panels (a)–(c) show the spatial maps corresponding to the first three tensor components obtained from CP decomposition, capturing dominant spatial–molecular patterns across the tissue. Panels (d)–(f) present the derived network representations, including the thresholded binary similarity network, node strength map, and local entropy map.

**Figure 4:**
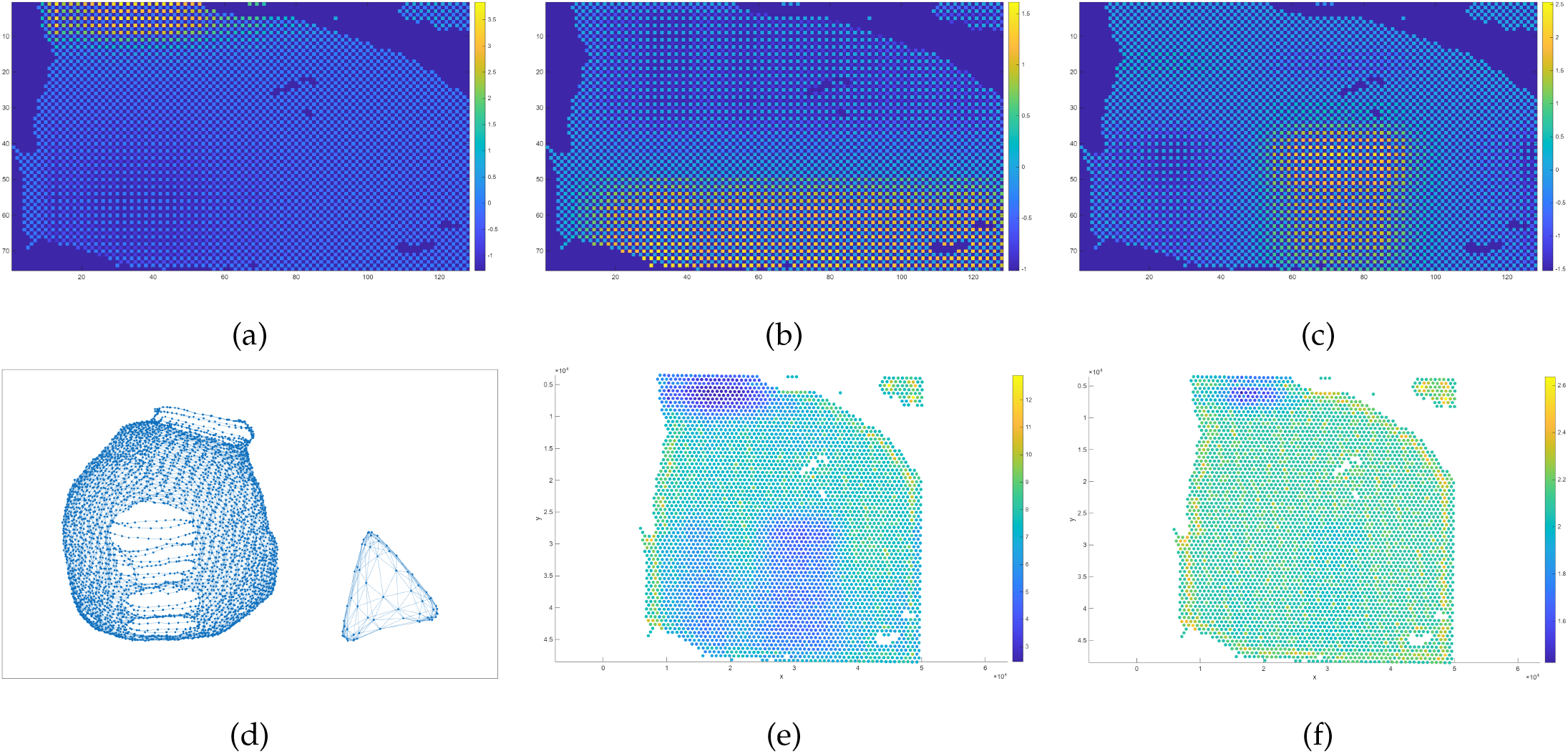
Visualisation of results for Patient 1. Panels show (a)–(c) spatial maps of tensor components 1–3, (d) thresholded binary similarity network, (e) node strength map, and (f) local entropy map.

**Figure 5:**
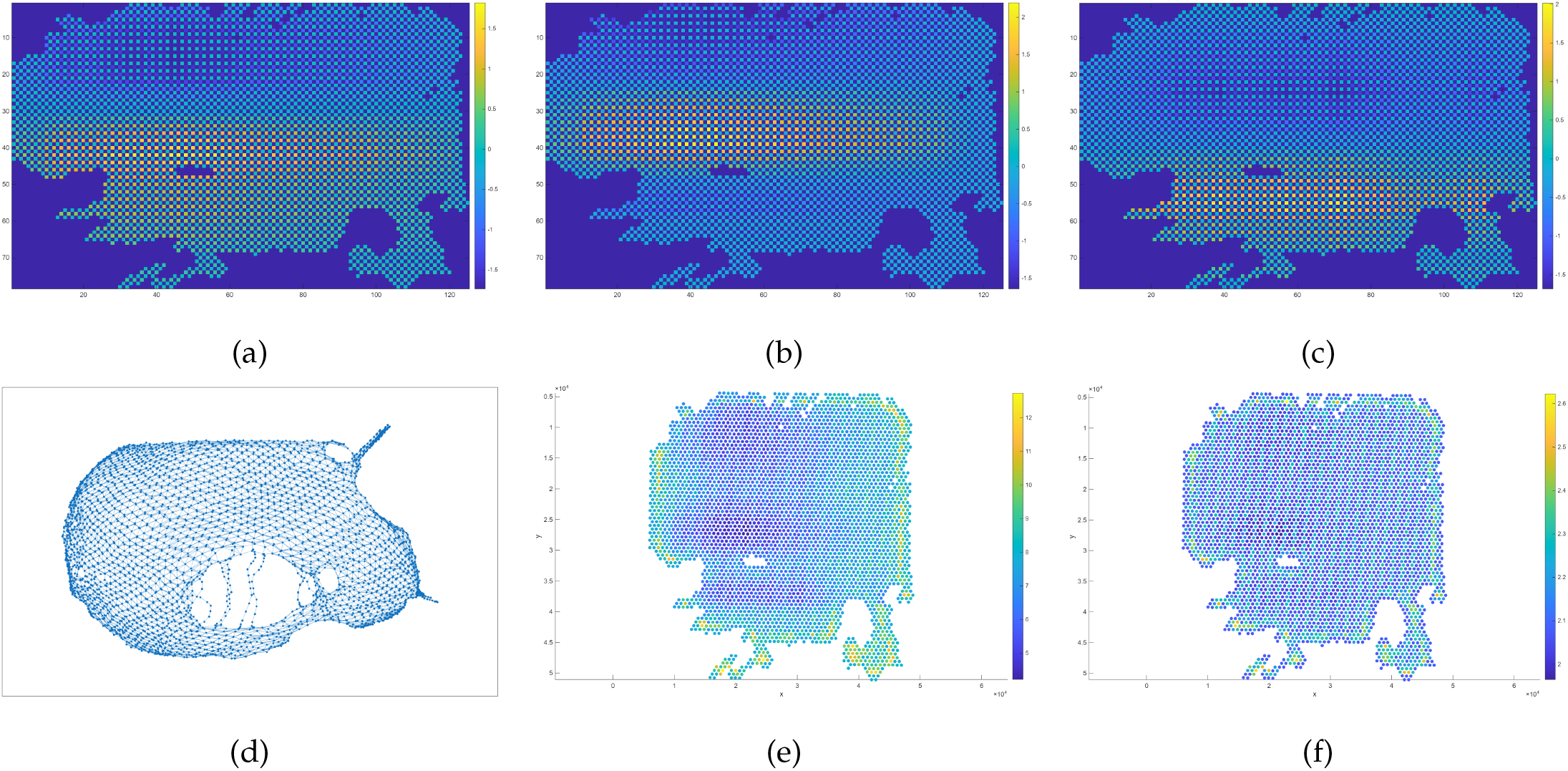
Visualisation of results for Patient 2. Panels show (a)–(c) spatial maps of tensor components 1–3, (d) thresholded binary similarity network, (e) node strength map, and (f) local entropy map.

**Figure 6:**
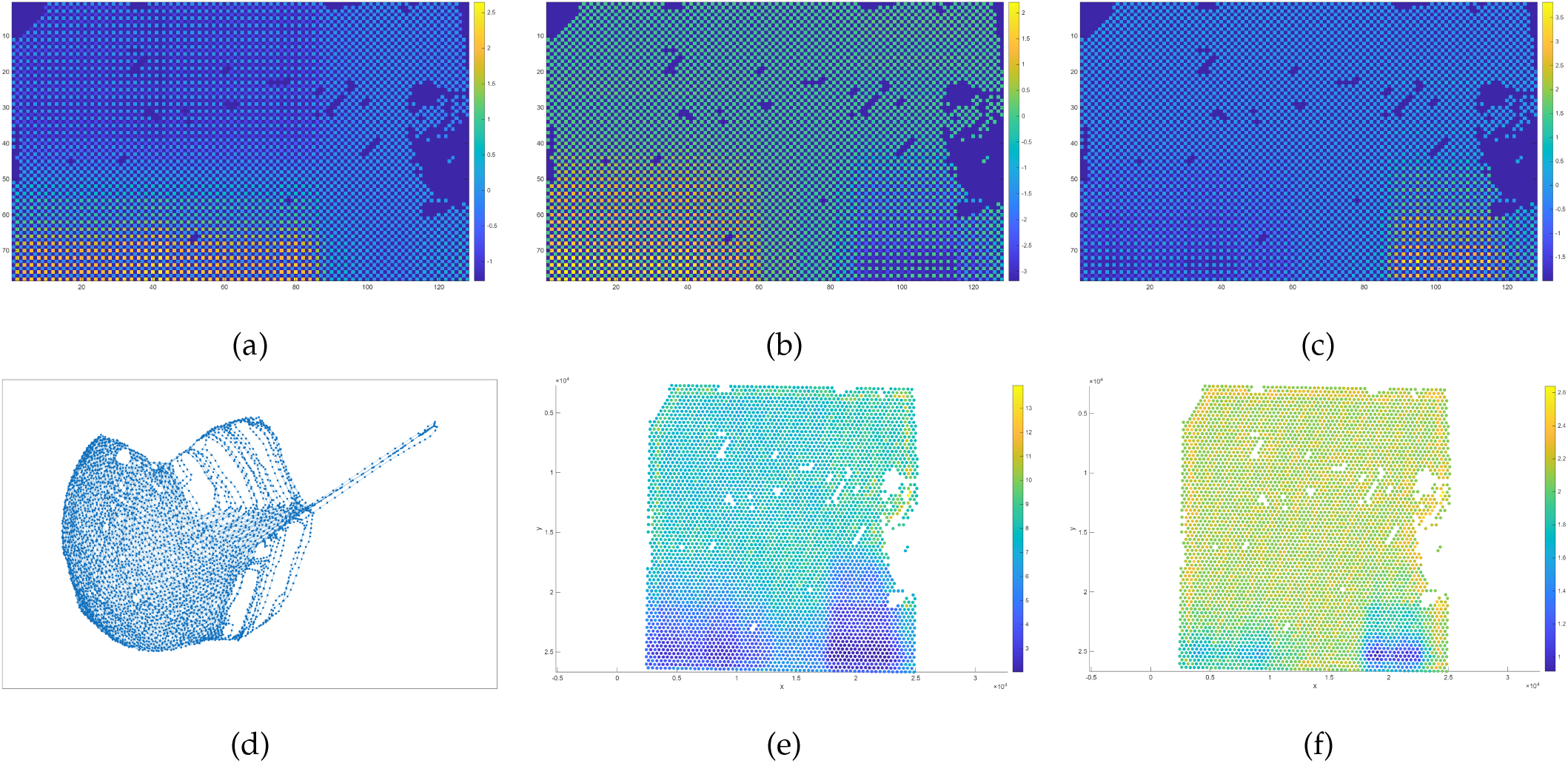
Visualisation of results for Patient 3. Panels show (a)–(c) spatial maps of tensor components 1–3, (d) thresholded binary similarity network, (e) node strength map, and (f) local entropy map.

**Figure 7:**
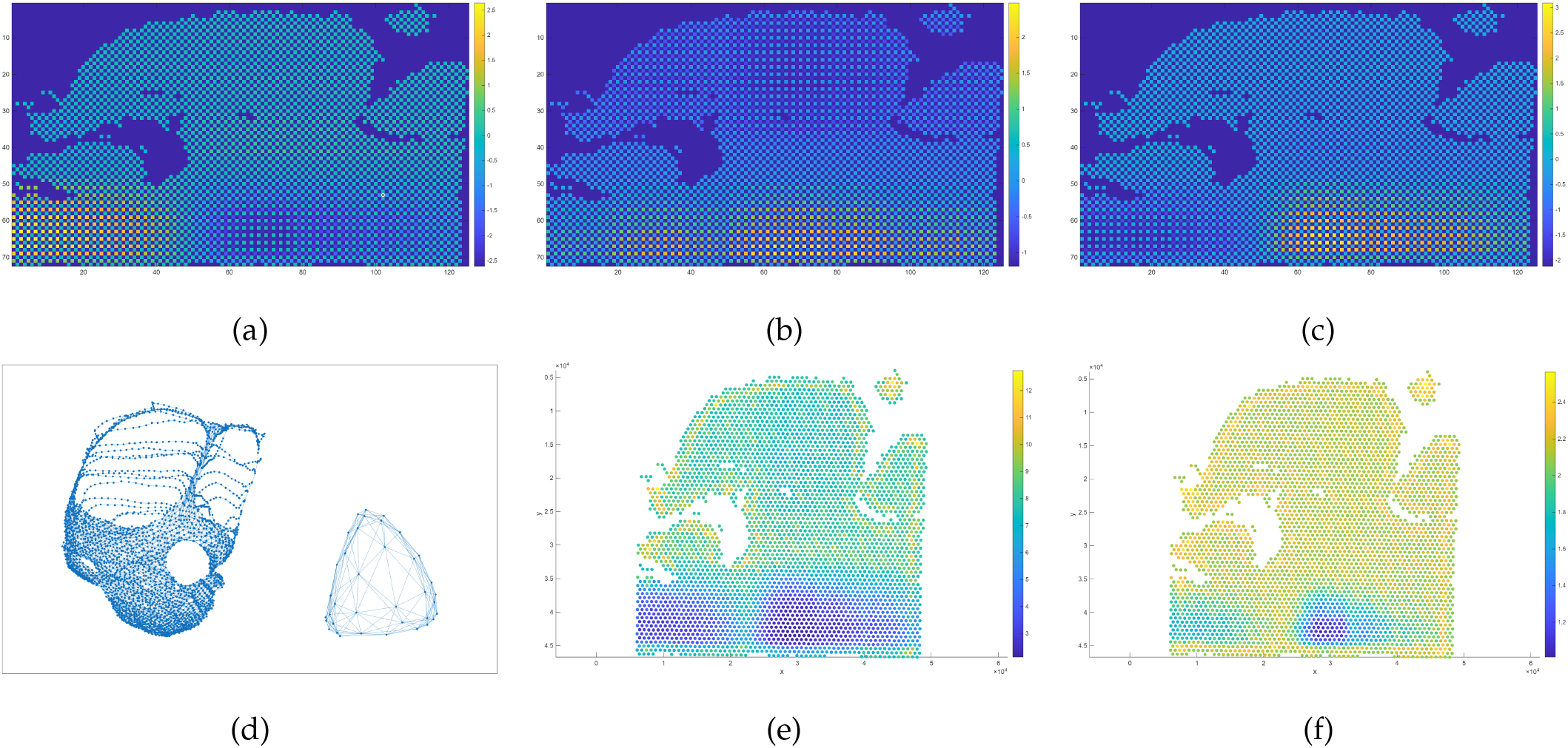
Visualisation of results for Patient 4. Panels show (a)–(c) spatial maps of tensor components 1–3, (d) thresholded binary similarity network, (e) node strength map, and (f) local entropy map.

The embedding-permutation framework results are shown in Figures 8, 9, 10, and 11. For each patient, panels (a)–(c) display the corresponding network representations obtained after random permutation of the embedding across spatial locations. The spatial maps of the tensor components remain unchanged as they are derived from the original tensor decomposition.

**Figure 8:**
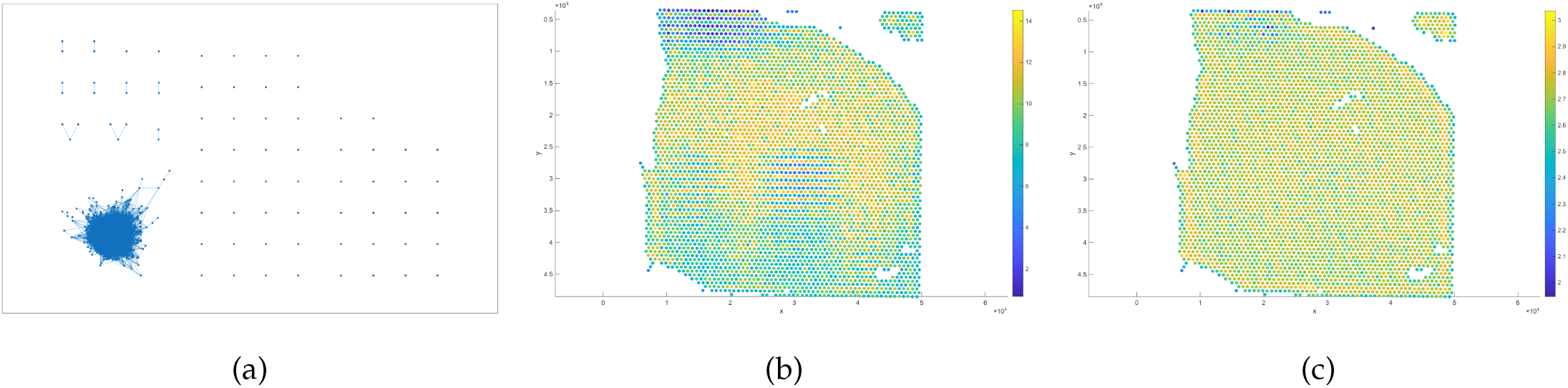
Visualisation of results for an embedding-permuted configuration of Patient 1. Panels show (a) thresholded binary similarity network, (b) node strength map, and (c) local entropy map..

**Figure 9:**
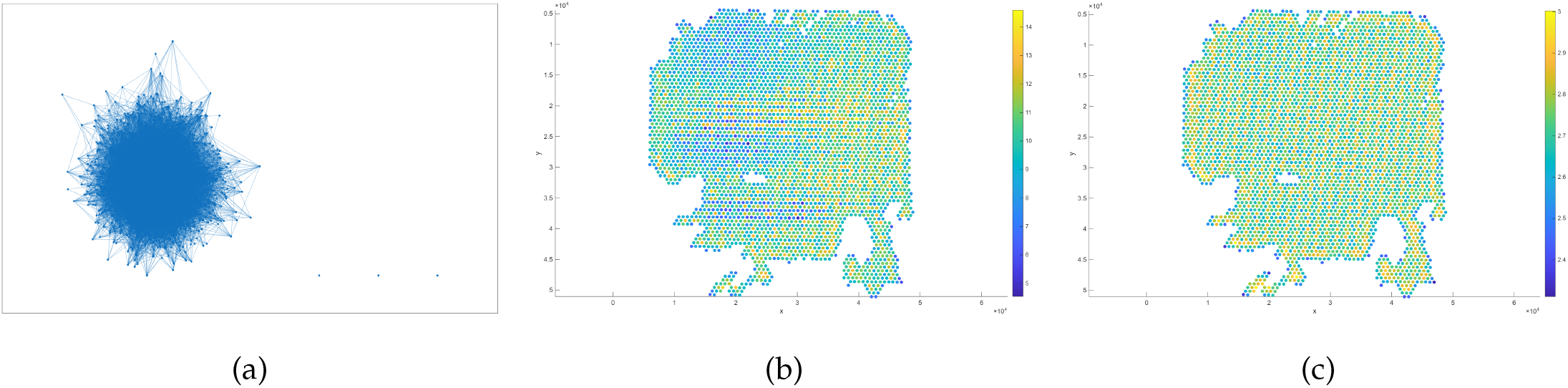
Visualisation of results for an embedding-permuted configuration of Patient 2. Panels show (a) thresholded binary similarity network, (b) node strength map, and (c) local entropy map.

**Figure 10:**
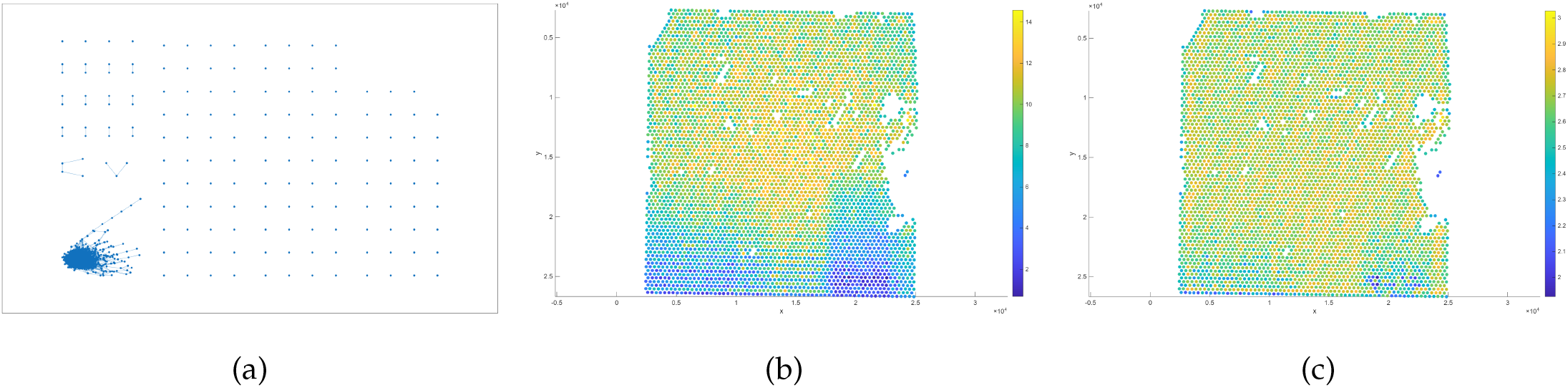
Visualisation of results for an embedding-permuted configuration of Patient 3. Panels show (a) thresholded binary similarity network, (b) node strength map, (c) local entropy map.

**Figure 11:**
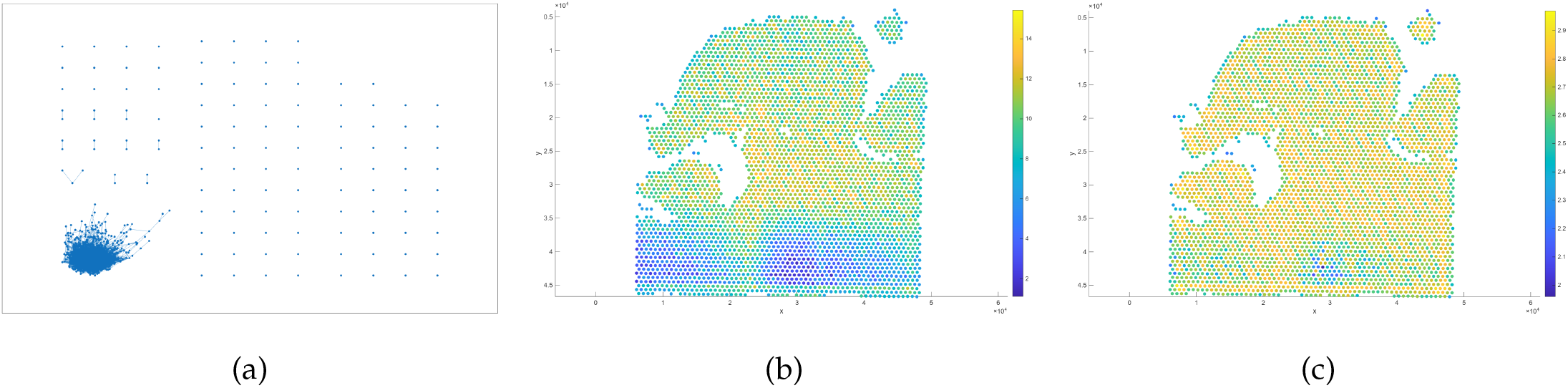
Visualisation of results for an embedding-permuted configuration of Patient 4. Panels show (a) thresholded binary similarity network, (b) node strength map, and (c) local entropy map.

## 4 Discussion

### 4.1 Tensor-Derived Representations of Spatial Transcriptomics Data

The application of the tensor-derived similarity network framework across four CRC patients demonstrates that high-dimensional spatial transcriptomics data can be effectively reduced into compact latent representations while preserving spatial structure. As shown in Table 1, despite variability in the number of spatial spots across patients, the tensor construction and CP decomposition yield consistent embedding dimensions (*N ×* 5), enabling comparable downstream network analysis.

The use of a rank-*K* = 3 CP model provides a low-dimensional representation that captures dominant spatial–molecular patterns while maintaining computational efficiency. The consistency of tensor sizes and embedding dimensions across patients suggests that the proposed framework is robust to variations in tissue size and sampling density.

### 4.2 Characteristics of Similarity Networks in Real Spatial Configurations

The global network measures in Table 2 indicate that the tensor-derived similarity networks are sparse across all patients, as reflected by low values of global similarity *R*_global_ and network density *D*. This sparsity suggests that strong similarity relationships are limited to specific subsets of spatial spots rather than being uniformly distributed across the tissue.

Despite this sparsity, the largest connected component contains the majority of nodes in each patient, indicating that the networks remain globally connected. This combination of sparse connectivity and large connected components suggests the presence of structured but selective spatial relationships within the tissue.

The global heterogeneity measure *HT* varies across patients, with lower values observed in Patient 2 and higher values in Patients 3 and 4. This variation reflects differences in the distribution of local similarity patterns, indicating that spatial organisation is not uniform across patients and may capture differences in underlying tumour architecture.

### 4.3 Behaviour of Networks under Random Spatial Organisation

The embedding-permutation framework provides a reference for understanding how network measures behave in the absence of spatial organisation. As shown in Table 3, the permuted configurations consistently exhibit higher values of global similarity, network density, and spatial heterogeneity compared to the real data.

The increase in *R*_global_ and *D* under permutation indicates that random reassignment of embedding vectors leads to a more uniformly connected network. This suggests that the structured spatial arrangement in the real data constrains connectivity, resulting in sparser networks.

Similarly, the higher values of *HT* in the permuted configurations indicate increased variability in local similarity distributions when spatial alignment is removed. The small standard deviations across permutations demonstrate that these effects are stable and reproducible, indicating that the observed behaviour is inherent to the randomisation process rather than due to variability in individual permutations.

### 4.4 Comparison between Real and Randomised Configurations

The comparison in Table 4 highlights systematic differences between real and embedding-permuted configurations across all patients. In the real data, lower values of *R*_global_ and *D* indicate more selective and spatially constrained similarity relationships, whereas the higher values observed under permutation reflect the loss of spatial structure.

The contrast in global heterogeneity *HT* further emphasises this difference. The real configurations exhibit lower heterogeneity compared to the permuted cases, indicating that spatial organisation imposes structure on local similarity patterns. In contrast, randomised configurations produce more uniform and less structured distributions of similarity, resulting in higher entropy.

Across all patients, the consistent separation between real and permuted metrics suggests that the tensor-derived embeddings encode spatially organised information that is not preserved under random permutation. This indicates that the observed network structures are influenced by spatial organisation rather than arising solely from the distribution of embedding features.

### 4.5 Implications for Spatial Tumour Heterogeneity

Taken together, the results indicate that the tensor-derived similarity network framework captures spatially structured patterns in CRC tissue. The sparsity of real networks, combined with their large connected components and lower heterogeneity relative to random configurations, suggests that tumour organisation is characterised by structured yet non-uniform spatial relationships.

The differences observed between patients further indicate that spatial tumour heterogeneity varies across specimens, as reflected in the variability of network measures. The proposed framework therefore provides a quantitative approach for characterising spatial organisation and comparing tumour structure across patients.

### 4.6 Interpretation of Tensor-Derived Spatial Patterns

Figures 4–7 illustrate the tensor-derived similarity network results for the four CRC patients. Panels (a)–(c) in each figure correspond to the spatial distributions of the first three tensor components, revealing dominant spatial–molecular patterns across the tissue. Across all patients, these components exhibit smooth spatial gradients and localised regions of elevated intensity, indicating that the tensor decomposition captures structured variation in gene expression aligned with tissue morphology.

In Patient 1 (Figure 4), the tensor components show relatively smooth transitions across the spatial grid, with distinct regions of higher activation emerging in specific areas. The corresponding similarity network (panel (d)) forms a well-connected structure with spatial coherence, while the node strength map (panel (e)) highlights regions with stronger local similarity. The entropy map (panel (f)) indicates moderate variability, with heterogeneous regions concentrated in specific spatial zones.

For Patient 2 (Figure 5), the tensor components display more pronounced spatial localisation, with identifiable regions of higher intensity embedded within a broader background. The similarity network remains connected but exhibits variations in local connectivity patterns. The node strength map reveals a more heterogeneous distribution compared to Patient 1, and the entropy map indicates spatial variability distributed across the tissue.

In Patient 3 (Figure 6), the tensor components show stronger regional contrast, with distinct high-intensity areas. The similarity network structure appears slightly less uniform, reflecting variability in spatial relationships between spots. The node strength and entropy maps demonstrate increased heterogeneity, with more pronounced spatial variation across the tissue.

In Patient 4 (Figure 7), the tensor components again exhibit structured spatial patterns, but with a different organisation compared to the other patients. The similarity network shows clear spatial connectivity patterns, while the node strength map highlights regions of strong local similarity. The entropy map indicates relatively high spatial variability, suggesting a more complex spatial organisation.

### 4.7 Interpretation of Embedding-Permuted Configurations

Figures 8–11 present the corresponding results obtained from the embedding-permutation framework, where the spatial alignment of tensor-derived embeddings is randomised. In contrast to the real configurations, the network structures in these figures reflect the absence of spatial organisation.

For Patient 1 (Figure 8), the thresholded similarity network (panel (a)) exhibits a highly centralised structure with connections concentrated in a limited region rather than following the spatial layout of the tissue. The node strength map (panel (b)) appears more uniform across the spatial grid, and the entropy map (panel (c)) shows elevated and relatively homogeneous values, indicating increased randomness in local similarity distributions.

A similar pattern is observed for Patient 2 (Figure 9), where the similarity network lacks the spatial coherence seen in the real data. The node strength distribution becomes more evenly spread, and the entropy map shows reduced spatial structure, reflecting the disruption of spatial relationships through permutation.

For Patient 3 (Figure 10), the permuted network again exhibits a centralised and less spatially structured connectivity pattern. The node strength and entropy maps demonstrate a more uniform distribution compared to the corresponding real configuration, indicating that the variability observed in the original data is largely diminished under randomisation.

In Patient 4 (Figure 11), the embedding-permuted configuration produces a network with similar characteristics, including reduced spatial coherence and increased uniformity in node strength and entropy. The entropy map shows consistently high values across the tissue, reflecting the loss of structured spatial heterogeneity.

### 4.8 Comparison Between Real and Permuted Configurations

Comparing Figures 4–7 with Figures 8–11 highlights clear differences between real and randomised spatial configurations. In the real data, the similarity networks exhibit spatially coherent structures aligned with tissue organisation, and the node strength and entropy maps reveal localised patterns of similarity and heterogeneity. In contrast, the embedding-permuted configurations produce networks that lack spatial coherence, with more uniform node strength distributions and elevated entropy across the tissue.

These differences indicate that the tensor-derived embeddings capture spatially organised patterns that are not preserved under random permutation. The observed contrast between structured spatial maps in the real data and homogenised patterns in the permuted configurations reflects the role of spatial organisation in shaping the similarity network topology.

### 4.9 Clinical Relevance and Potential Applications

The tensor-derived similarity network framework provides a quantitative representation of spatial organisation in CRC tissue, with potential relevance for clinical interpretation and decision-making. By transforming high-dimensional spatial transcriptomics data into structured network representations, the proposed approach enables the characterisation of tumour architecture beyond conventional histopathological assessment.

The observed differences in global similarity, network density, and spatial heterogeneity across patients suggest that the framework captures variations in tumour organisation that may reflect underlying biological processes. In a clinical context, such differences could be associated with tumour aggressiveness, invasion patterns, or microenvironmental complexity. For example, lower network density and similarity in real spatial configurations indicate selective and spatially constrained interactions, which may correspond to compartmentalised tumour regions or heterogeneous cellular niches.

The spatial heterogeneity measure *HT* provides a compact summary of local variability across the tissue. Higher values of *HT* may indicate more complex tumour architecture, potentially associated with increased biological diversity within the tumour microenvironment. Conversely, lower values may reflect more homogeneous tissue organisation. These properties suggest that network-based metrics could serve as quantitative biomarkers for assessing spatial tumour heterogeneity.

Furthermore, the embedding-permutation framework demonstrates that the observed network structures are not solely determined by the distribution of molecular features but depend on their spatial arrangement. This highlights the importance of spatial context in interpreting transcriptomic data, which is increasingly recognised as critical in precision oncology.

From a translational perspective, the proposed framework could support several applications. First, it may aid in the identification of spatial biomarkers by highlighting regions of high node strength or entropy, which could correspond to biologically distinct tumour subregions. Second, it could be integrated with imaging and histopathology to provide multimodal characterisation of tumour structure. Third, the quantitative measures derived from similarity networks could be incorporated into predictive models for patient stratification, prognosis, or treatment response.

Overall, the tensor-derived similarity network approach offers a systematic method for capturing spatial organisation in tumour tissue, providing a foundation for future studies linking spatial transcriptomic patterns to clinical outcomes.

### 4.10 Limitations and Future Work

Several limitations of the present study should be acknowledged. First, the analysis was conducted on a relatively small cohort of four CRC patients, which limits the generalisability of the findings. Although the framework demonstrates consistent behaviour across patients, larger and more diverse datasets are required to validate its robustness and clinical relevance.

Second, the tensor decomposition was performed using a fixed rank (*K* = 3), which may not fully capture the complexity of spatial–molecular patterns in all tissues. While this choice provides a compact and interpretable representation, the selection of tensor rank remains an important modelling consideration [36, 50, 51, 52]. Future work could explore data-driven approaches for rank selection or adaptive decomposition strategies.

Third, the similarity network construction relies on a Gaussian kernel and a spatial *k*-nearest neighbour graph, which introduce hyperparameters such as the kernel scale and neighbourhood size. Although these were chosen to provide stable results, the sensitivity of the framework to these parameters warrants further investigation.

Fourth, the embedding-permutation framework provides a reference for random spatial organisation but does not constitute a formal statistical testing procedure in the current study. While mean and standard deviation of network measures are reported, future work could incorporate more rigorous statistical inference, including hypothesis testing or confidence intervals, to quantify the significance of observed differences.

From a biological perspective, the study focuses on computational characterisation of spatial patterns without direct integration of histopathological annotations or clinical outcomes. Consequently, the biological interpretation of the network measures remains indirect. Future studies should aim to link tensor-derived network features with tumour subtypes, microenvironmental composition, and patient prognosis.

In addition, the current framework considers a single modality of spatial transcriptomics data. Extensions to multimodal data, such as integrating spatial proteomics, imaging, or clinical variables, could provide a more comprehensive representation of tumour organisation.

Finally, the computational pipeline, while efficient, could be further optimised for large-scale datasets and extended to include advanced machine learning approaches, such as graph neural networks or deep generative models, to enhance representation learning and predictive capability.

Overall, these limitations highlight opportunities for methodological refinement, biological validation, and clinical translation in future work.

## 5 Conclusion

This study introduced a tensor-derived similarity network framework for analysing spatial transcriptomics data in CRC. By integrating tensor decomposition with network-based analysis, the approach provides a structured representation of spatial–molecular patterns within tumour tissue. The resulting similarity networks enable quantification of spatial organisation through global measures of similarity, density, and heterogeneity.

Across all patients, the framework revealed sparse yet structured connectivity patterns, indicating that spatial relationships between tissue regions are selective rather than uniformly distributed. The embedding-permutation framework further demonstrated that these structures are influenced by spatial organisation, as randomised configurations consistently produced more densely connected and more heterogeneous networks.

The proposed methodology offers a systematic approach for characterising spatial tumour heterogeneity and provides a foundation for further investigation of spatial patterns in cancer. Future work will focus on extending the framework to larger cohorts, integrating multimodal data, and linking network-derived measures to biological and clinical outcomes. Overall, this study highlights the potential of tensor-based network modelling as a tool for advancing the analysis of spatially resolved molecular data.

